# Evaluating Fairness and Generalizability of Alzheimer’s Disease Diagnosis Models Trained on Racially Imbalanced Datasets

**DOI:** 10.1101/2025.09.30.678854

**Authors:** Neha Goud Baddam, Bizhan Alipour Pijani, Melissa Petersen, Serdar Bozdag

**Affiliations:** Department of Computer Science and Engineering, University of North Texas, Denton, Texas, USA; Center for Computational Life Sciences, University of North Texas, Denton, Texas, USA; BioDiscovery Institute, University of North Texas, Denton, Texas, USA; Institute for Translational Research, University of North Texas Health Science Center, Fort Worth, Texas, USA; Department of Family Medicine, University of North Texas Health Science Center, Fort Worth, Texas, USA; Department of Mathematics, University of North Texas, Denton, Texas, USA

## Abstract

Alzheimer’s disease (AD) is a major global health concern, expected to affect 12.7 million Americans by 2050. Machine learning (ML) algorithms have been developed for AD diagnosis and progression prediction, but the lack of racial/ethnic diversity in clinical datasets raises concerns about their generalizability across demographic groups, particularly underrepresented populations. Studies show ML algorithms inherit biases from data, leading to biased AD predictions. This study investigates the fairness of ML models in AD diagnosis. We hypothesize that models trained on a single racial/ethnic group perform well within that group but poorly in others. We employ feature selection and model training techniques to improve fairness. Our findings support our hypothesis that ML models trained on one group underperform on others. We also demonstrated that applying fairness techniques to ML models reduces their bias. This study highlights the need for racial/ethnic diversity in datasets and fair models for AD prediction.

## 1 BACKGROUND

Alzheimer’s disease (AD) is a serious neurodegenerative disease that poses a global medical threat. The Alzheimer’s Association estimates that 6.9 million Americans had Alzheimer’s in 2024, and by 2050, that number will rise to 12.7 million. Recently, computational methods have been developed to diagnose AD and track its progression. These models are promising due to their ability to handle large data, perform complex computations, and make impartial decisions. However, studies like [1] argue that this assumption of impartiality is questionable, as ML algorithms often inherit and amplify biases in their training data [2]. Fairness in AI refers to the absence of bias or discrimination against individuals or groups based on *sensitive attributes*, such as race, and features that reflect socio-cultural aspects. Biases may stem from historical inequities in healthcare when certain groups (e.g., minority racial/ethnic groups) have been affected. For example, research has shown variability in healthcare access for certain racial/ethnic groups [3, 4]. Missing or incomplete data further affects predictions, as some populations are underrepresented in training data [5]. AD datasets often lack representation from specific racial/ethnic groups, raising concerns about bias [6]. For instance, studies [7] note that the widely used ADNI dataset has oversampled White/Caucasian individuals.

To address fairness in AI, researchers propose various strategies. *Pre-processing methods* modify input data before training by re-sampling [8], re-weighting [9], or adjusting features to balance group representation. Some studies apply fairness-aware feature selection. For example, [11] selects features minimally correlated with sensitive attributes, while tools like FairML [12] rank features for fair predictions. *In-processing methods* embed fairness directly into the learning algorithm, modifying the loss function to penalize bias [13] or applying adversarial training to reduce the model’s reliance on protected attributes [14]. *Post-processing methods* adjust predictions to ensure fairness by re-labeling outputs [15] or perturbing non-sensitive attributes [16].

Fairness in ML is measured through metrics, broadly classified into group fairness (treating groups equally) and individual fairness (treating similar individuals similarly) [17]. Most metrics are built for binary classification with binary sensitive attributes [18], but healthcare often involves multiple outcomes, such as distinguishing between normal cognition, mild cognitive impairment (MCI), and AD. Despite progress in AD prediction, three limitations remain:

### Dataset limitations

Models trained on non-diverse datasets may not generalize well across racial/ethnic groups.

### Model limitations

Though fairness-aware methods exist, they are rarely applied to healthcare, particularly for AD diagnosis.

### Metric limitations

Some fairness metrics have been adapted for multi-class tasks and multi-category sensitive attributes [19], but lack thorough testing in healthcare.

Motivated by these gaps, our study explores the fairness of ML models in diagnosing AD, focusing on racial/ethnic bias. We hypothesize that models trained on one racial/ethnic group perform best on that group but may not generalize to others. For this work, we utilized *Health &Aging Brain Study: Health Disparities* (HABS-HD) dataset [20], which includes African American and Mexican American participants, groups often underrepresented in AD studies, making it suitable for evaluating fairness. To test our hypothesis, we train models on one racial/ethnic group and evaluate on others. Then, we trained a model on data from all racial/ethnic groups and assessed performance for each racial/ethnic group to see if disparities persist. To improve fairness, we applied both pre-processing and in-processing techniques. Our work expands fairness research by addressing multi-class classification and non-binary sensitive attributes common in healthcare.

### Our main contributions

1. We assessed the generalizability of ML models trained on one racial/ethnic group when tested on others.
2. We evaluated whether diverse training data improves fairness across racial/ethnic groups.
3. We implemented and evaluated fairness-aware feature selection and in-processing strategies to mitigate racial/ethnic bias.
4. We extended fairness metrics for multi-class classification with multi-category sensitive attributes, reflecting real-world applications.

Our experiments using HABS-HD show that when ML models trained on a single racial/ethnic group, they exhibit racial/ethnic bias. Applying fairness-aware techniques and utilizing more diverse training data helps mitigate this bias and improves model performance across groups.

## 2 RESULTS

In this section, we summarize the evaluation of our experimental approach for different scenarios explained in Section 4.3. We evaluated the performance of ML models for AD diagnosis prediction, where the class labels were AD, MCI, and CN. To achieve robust results, we considered several ML algorithms and chose RF as it demonstrated the best predictive performance (see Section 4.3). Additionally, we incorporated fairness-aware evaluation metrics such as DP, EOd, and EOp (see Section 4.5 for their definitions). Lower values of DP, EOd, and EOp indicate better fairness (i.e., smaller disparities across racial/ethnic groups).

As shown in **Table 2**, we conducted a cross-racial/ethnic evaluation by training the model on data from each racial/ethnic group and testing it on all the others. This analysis aimed to explore whether models trained exclusively on one racial/ethnic group would generalize poorly to other groups. In all three cases (i.e., training group AA, MA, and NHW), this hypothesis was statistically supported as the models achieved significantly better performance on their own group compared to the other two groups (p-values ≤4.55×10^−29^ for macro F1 and p-values ≤5.83 ×10^−29^ for balanced accuracy). For instance, the model trained on the MA group achieved a macro F1 score of 0.957 on MA but dropped to 0.560 and 0.676 on AA and NHW, respectively. Statistical significance between groups was evaluated using Welch’s t-test. Similarly, the NHW-trained model performed best on NHW (macro F1 score: 0.966) and showed reduced scores on AA (0.873) and MA (0.753). The AA-trained model also followed this pattern, achieving the highest performance on AA (macro F1: 0.904) compared to MA (0.659) and NHW (0.760). These results indicate that models trained on data from a single racial/ethnic group consistently achieve strong performance within that group but show reduced performance when applied to other groups, thereby supporting our hypothesis. These discrepancies highlight the importance of inclusive training data. The fairness metrics DP, EOd, and EOp further emphasize disparities across racial/ethnic groups (See **Table 3**), showing that models trained on one racial/ethnic group not only perform poorly on the other racial/ethnic groups but also exhibit higher levels of bias.

**Table 1:** Mean and standard deviation of balanced accuracy and Macro F1 score of the RF models (features from Lasso), trained on data from one racial group and tested on each racial group separately (NHW: Non-Hispanic White, AA: African American, MA: Mexican American). Bold values indicate the best performance for each model. Values are reported as mean ±standard deviation across 50 independent runs with different random seeds. ↑indicates higher values are better

| Training Group | Test Group | Balanced Accuracy ↑ | Macro F1 Score ↑ |
| --- | --- | --- | --- |
| AA | AA | **0.897±0.062** | **0.904±0.059** |
| | MA | 0.753±0.059 | 0.659±0.095 |
| | NHW | 0.804±0.080 | 0.760±0.108 |
| MA | AA | 0.657±0.123 | 0.560±0.181 |
|  | MA | **0.949±0.035** | **0.957±0.030** |
| | NHW | 0.718±0.078 | 0.676±0.105 |
| NHW | AA | 0.914±0.052 | 0.873±0.099 |
| | MA | 0.791±0.051 | 0.753±0.070 |
|  | NHW | **0.958±0.032** | **0.966±0.027** |

**Table 2:** Fairness measures of models trained on data from a single racial group and tested on other racial groups (NHW: Non-Hispanic White, AA: African American, MA: Mexican American). Values are reported as mean ±standard deviation across 50 independent runs with different random seeds. ↓indicates lower values are better.

| Training Group | Demographic Parity (DP) ↓ | Equalized Odds Difference (EOd) ↓ | Equal Opportunity (EOp) ↓ |
| --- | --- | --- | --- |
| AA | 0.112±0.027 | 0.703±0.250 | 0.609±0.223 |
| MA | 0.300±0.286 | 0.977±0.510 | 0.627±0.201 |
| NHW | 0.144±0.092 | 0.550±0.219 | 0.434±0.156 |

**Table 3:** Comparison of baseline (features from SelectbySinglefeaturePerformance), pre-processing (features selected using multiple feature selection methods), and combined pre- and in-processing approaches (integrating feature selection with model-based weighting) on various performance and fairness metrics across racial test groups (NHW: Non-Hispanic White, AA: African American, MA: Mexican American). Values are reported as mean *±*standard deviation across 50 independent runs with different random seeds. ↑indicates higher values are better, while ↓indicates lower values are better.

| Metrics | Test Group | Baseline | Pre-processing | Pre- & In-processing |
| --- | --- | --- | --- | --- |
| Balanced Accuracy ↑ | AA | **0.944±0.034** | 0.941±0.039 | 0.943±0.038 |
| | MA | 0.903±0.043 | 0.912±0.042 | **0.919±0.041** |
| | NHW | 0.966±0.024 | **0.972±0.023** | 0.968±0.022 |
| Macro F1 Score ↑ | AA | **0.950±0.033** | 0.946±0.038 | 0.950±0.036 |
| | MA | 0.914±0.039 | 0.922±0.038 | **0.927±0.038** |
| | NHW | 0.971±0.022 | **0.976±0.021** | 0.973±0.019 |
| Demographic Parity (DP) ↓ | | 0.069±0.008 | **0.067±0.007** | 0.068±0.008 |
| Equalized Odds Difference (EOd) ↓ | | 0.244±0.125 | 0.223±0.119 | **0.215±0.111** |
| Equal Opportunity (EOp) ↓ | | 0.213±0.120 | 0.194±0.115 | **0.180±0.108** |

To evaluate whether training on a racially/ethnically diverse dataset results in equitable performance across racial/ethnic groups (See Section 4.3), we evaluated a model trained on samples from all three racial/ethnic groups and tested on each racial/ethnic group (baseline column in **Table 4**). Compared to the results observed in the cross-racial/ethnic evaluation (See **Table 2** and **Table 3**), we observed that the baseline model performed better in terms of fairness. This is likely due to the broader representation in the training data, which enables the model to generalize better across racial/ethnic groups. The improvements in fairness were evident across all the fairness metrics, where the baseline model reduced DP to 0.069, EOd to 0.244, and EOp to 0.213, compared to much higher values observed in the cross-racial/ethnic evaluation (e.g., DP of 0.300, EOd of 0.977, and EOp of 0.627 for the MA-trained model).

**Table 4:** Top 10 most influential features selected by the LASSO feature selection method for each racial/ethnic group (AA, NHW, MA).

| Rank | AA | NHW | MA |
| --- | --- | --- | --- |
| 1 | CDR_Memory | CDR_Memory | ID_Language_Secondary_Other |
| 2 | 01_AB_FBB_LatParietal_SUVr | CDR_HomeandHobbies | CDR_Sum |
| 3 | CDR_CA_NoFunctions | CDX_SCC | 01_AB_FBB_LatTemporal_SUVr |
| 4 | IMH_Alzheimers | CDR_PC_Dressing | 01_AB_FBB_LatParietal_SUVr |
| 5 | MMSE_3 | SCC_YesNo | CDR_Global |
| 6 | BW_ALT | CDR_PC_EatingHabits | 03_FA_Splenium_of_corpus_callosum_left |
| 7 | GDS_Score_D | MMSE_13a | PED_1_D |
| 8 | AMNART_Er | CDR_Judgement | EDS_8 |
| 9 | DSB_1_T2 | PED_3_C | AMNART_2g |
| 10 | 01_L_isthmuscingulate_thickavg | SCC_54I | 03_AD_Cingulum_cingulate_gyrus_right |

In terms of predictive performance, the baseline method outperformed the cross-racial/ethnic models trained and tested on their own groups in two of three groups. For instance, the baseline method achieved a macro F1 score of 0.950 for the AA group, which was higher than the macro F1 score of the AA-trained model on the AA test samples (0.904). Despite these improvements, we observed some disparities in performance across racial/ethnic groups in the baseline model. For example, the macro F1 score for NHW individuals was 0.971, but it dropped to 0.950 for AA and 0.914 for the MA group, which suggests that the baseline model still suffers from fairness issues.

To address the observed disparities, we implemented both fairness-aware pre-processing and in-processing model training strategies. The results, summarized in **Table 4**, demonstrate improvements in all fairness metrics and predictive performance in some cases, compared to the baseline model. The inclusion of fairness interventions led to reductions in disparity as measured by DP, EOd, and EOp. For instance, EOp decreased from 0.213 in the baseline model to 0.197 after applying both pre-processing and in-processing techniques. We observed that both the balanced accuracy and macro F1 score dropped slightly in these models compared to the baseline model for the AA groups, while we observed a slight improvement in the predictive performance for the MA and NHW groups.

The findings from this study underscore the importance of equitable model performance, especially when deploying ML systems in settings where social equity is critical, such as healthcare, finance, and criminal justice. Our results demonstrate that fairness-aware feature selection and model training strategies can reduce bias without compromising overall model utility. Moreover, our fairness metric extensions to the multiclass setting provide a comprehensive toolkit for practitioners to diagnose and address inequality in diverse real-world applications. These results emphasize the need for continuous fairness auditing and inclusive datasets as fundamental steps in responsible AI development.

## 3 DISCUSSION

In this study, we evaluated ML models based on fairness and predictive performance under different scenarios. Specifically, for AD diagnosis, we examined how ML models perform when trained on samples from one racial/ethnic group compared to the models trained on a racially/ethnically diverse training data. We also evaluated to what extent fairness techniques, such as pre-processing and in-processing, affect model performance. Our results showed a consistent pattern that the models trained using data from only one racial/ethnic group tended to yield the highest predictive performance for that same group. However, their performance dropped significantly when applied to individuals from other racial/ethnic backgrounds. This indicates that the models often learned group-specific patterns that did not generalize well to other groups, leading to biased predictions. This trend was strongly evident for the AA, MA, and NHW training groups, with statistically significant performance drops when tested on out-of-group data. To further investigate and mitigate these disparities (See **Table 4**), we trained the baseline model using data from all three racial/ethnic groups. We then tested the model separately on each racial/ethnic group to assess whether its performance was balanced across all groups. Interestingly, we found that even though the training data was more diverse this time, the model still performed better on some groups than others. This shows that simply including data from different racial/ethnic groups is not enough to eliminate bias or ensure fairness in predictions. To address this issue more effectively, we implemented two fairness-aware strategies. First, we applied a pre-processing technique, aiming to reduce bias in the input features. Second, we used pre- and in-processing techniques during model training that explicitly encouraged equitable treatment across groups. These combined approaches helped the model focus more on generalizable and less biased patterns. We measured fairness using three established metrics, namely DP, EOd, and EOp, which we adapted to our multiclass classification setting. Our results showed improvements across all fairness measures (DP, EOd, and EOp).

Beyond the immediate findings, several broader insights emerge from our study. **Table 5** lists the top 10 selected features for each racial/ethnic group, ranked by their importance scores by LASSO. In general, CDR-based features emerge as the most important across all three groups, which is not surprising as clinical diagnoses are mainly based on CDR-based scores. Among NHW, CDR Memory and other CDR-based functional measures stand out, as this group tends to rely strongly on clinical severity indicators. In contrast, identity and neuroimaging-related features, such as ID Language Secondary Other and AB FBB PET SUVR measures, were more prominent in the MA group. Interestingly, neuroimaging features appear in both AA and MA groups, but not prominently in the NHW group. Their ranking also varies: for example, 01 AB FBB LatParietal SUVR is ranked second among AA and fourth among MA. This suggests that while neuroimaging and cognitive measures are broadly relevant, their relative importance differs across populations. The selected features were further grouped into broader categories, following the HABS-HD dictionary. For example, CDR Memory, CDR Global, and CDR Sum were combined into the CDR category. **Table 6** presents the percentage distribution of features in each racial/ethnic group after applying this categorization. Notably, CDR-based features (Global/Clinical Severity) remain the most important across all groups, accounting for 20%of features in AA, 50%in NHW, and 20%in MA. Other feature groups also appear across populations, including AMNART (American National Adult Reading Test), SEVLT (Spanish English Version Language Test), MMSE (Mini-Mental State Examination), neuroimaging-derived measures, and PED (Physician’s Estimate of Duration) related variables, though their relative contributions vary across racial/ethnic groups. To further investigate feature representation patterns, we analyzed the distribution of feature families among the top 100 selected features for each racial/ethnic group. **Table 7** summarizes the five feature families with the highest representation across groups. Neuroimaging-derived features appear most frequently across all groups, accounting for 16%, 22%, and 29%of the top features in AA, NHW, and MA groups, respectively. CDR-based clinical severity measures are also consistently represented across groups, contributing 11%in AA, 12%in NHW, and 9%in MA. Cognitive assessment measures such as AMNART and SEVLT show varying levels of importance across populations, with AMNART appearing more frequently in MA (13%) compared to NHW (7%) and AA (5%). Similarly, SEVLT-based features contribute 9%of the selected features in AA but appear less frequently in NHW and MA (4%each). PED-related variables show moderate representation across groups, ranging from 3%in AA to 7%in NHW. These patterns suggest that while neuroimaging and clinical severity measures are broadly important for AD diagnosis prediction, the relative contribution of cognitive and functional assessment measures differs across racial/ethnic groups.

**Table 5:**
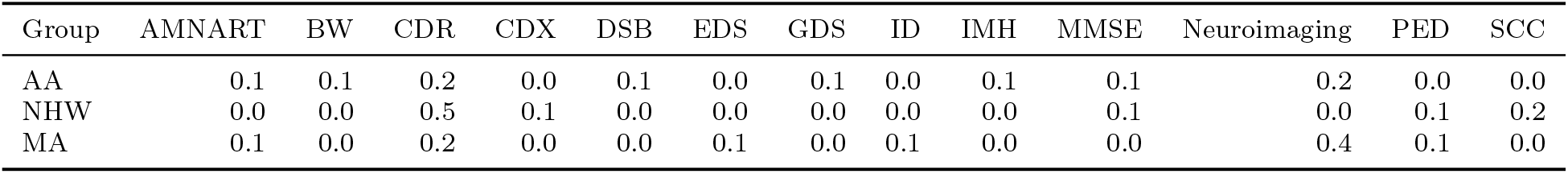
Percentage distribution of feature categories among the top 10 LASSO-selected features for each racial/ethnic group (AA, NHW, MA). These features were grouped into broader clinical and biological categories based on the HABS-HD dictionary (AMNART= American National Adult Reading Test, BW = Bloodwork, CDR = Clinical Dementia Rating, CDX = Clinical Diagnosis, DSB = Wechsler Memory Scale Third Edition Digit Span (Backward), EDS = Everyday Discrimination Scale, GDS = Geriatric Depression Scale, ID= Language-based features, IMH = Objective Medical History/Diagnosis, MMSE = Mini Mental State Examination, Neuroimaging = Imaging-derived biomarkers, PED = Physician’s Estimate of Duration of Dementia, SCC = Subjective Cognitive Complaints).

**Table 6:** Top 5 feature families with the highest representation among the top 100 features selected by LASSO for each racial/ethnic group (AA, NHW, MA). These features were grouped into broader clinical and biological categories based on the HABS-HD dictionary (Neuroimaging = Imaging-derived biomarkers, CDR = Clinical Dementia Rating, AMNART= American National Adult Reading Test, SEVLT = Spanish English Verbal Learning Test, PED = Physician’s Estimate of Duration of Dementia).

| Group | Neuroimaging | CDR | AMNART | SEVLT | PED |
| --- | --- | --- | --- | --- | --- |
| AA | 0.16 | 0.11 | 0.05 | 0.09 | 0.03 |
| NHW | 0.22 | 0.12 | 0.07 | 0.04 | 0.07 |
| MA | 0.29 | 0.09 | 0.13 | 0.04 | 0.04 |

**Table 7:** Distribution of HABS-HD samples across racial/ethnic groups and diagnostic categories (Cognitively Normal, Mild Cognitive Impairment, and Alzheimer’s Disease).

| Racial Group | Cognitively Normal | Mild Cognitive Impairment | Alzheimer’s Disease | Total Samples |
| --- | --- | --- | --- | --- |
| Non-Hispanic White (NHW) | 584 | 72 | 58 | 714 |
| Mexican American (MA) | 729 | 111 | 54 | 894 |
| African American (AA) | 396 | 30 | 26 | 452 |
| Total Samples | 1709 | 213 | 138 | 2,060 |

While our findings are promising and point in a positive direction, it is important to acknowledge that our study has certain limitations. Although the techniques we used and the findings we observed may apply to other areas of healthcare, such as breast cancer prediction, they have not been tested in our study. Furthermore, our model treats race as a fixed, well-defined attribute. In reality, race is a socially constructed concept influenced by historical, cultural, and systemic factors. It often intersects with other determinants like socioeconomic status, education, geography, and healthcare access, all of which can impact both health outcomes and data quality, yet were not fully accounted for in our study. Another limitation is that we did not explore longitudinal prediction of disease progression or evaluate fairness over time. AD evolves gradually, and disparities may change as patients transition between disease stages. Investigating fairness in a longitudinal setting, where models repeatedly forecast future cognitive decline, would be critical to ensure that equity is maintained at successive time points and disease trajectories. As future work, we plan to address these limitations by expanding the dataset and investigating how intersectional factors such as socioeconomic status and healthcare access interact with race to impact both outcomes and fairness evaluations, and by extending our framework to longitudinal prediction so that fairness can be assessed and enforced dynamically as disease progression unfolds.

In conclusion, our analysis of AD prediction revealed that an ML model can exhibit bias toward specific individuals in the prediction task when trained on data from a single racial/ethnic group. Providing a diverse dataset to the model training mitigates this bias to some extent, but does not fully resolve it. We addressed this issue by introducing two fairness techniques that help reduce model bias. Our findings suggest that combining pre-processing and in-processing techniques yields increased model fairness. These results underscore that fairness is not merely a property of the training data but an ongoing design consideration that must be addressed through algorithmic choices and continuous evaluation. By embedding fairness as a core design principle and extending these methods to larger and more diverse datasets, future studies can help ensure that ML evolves from a technology that may inadvertently reinforce health inequities into one that actively promotes equitable and trustworthy AD diagnosis.

## 4 METHODS

### 4.1 Dataset and Data Pre-processing

HABS-HD dataset offers a diverse representation of racial/ethnic groups, specifically Non-Hispanic White (NHW), Mexican American (MA), and African American (AA) individuals. HABS-HD participants are enrolled utilizing a community-based participatory research (CBPR) approach. Participant visits occur at 24-month intervals and include a clinical interview, neuropsychological assessment, blood draw, functional examination, MRI, and PET (amyloid and tau) scans. See for detailed methods. Cognitive diagnosis was derived through a consensus review process overseen by dementia experts associated with the study. Briefly, those determined to be Cognitively Normal (CN) did not present with a cognitive complaint and exhibited cognitive test performance that was determined to fall within what would be expected given the participants’demographics (normative range). In addition, the Clinical Dementia Rating scale (CDR) sum of boxes (SOB) and indication of functioning were scored at 0, noting intact functional abilities. Those determined to fall into the Mild Cognitive Impairment (MCI) designation were those with a cognitive complaint (by self or other) and had at least one score on their cognitive testing that fell at or below 1.5 SD from the normative range. For this diagnostic determination, the CDR SOB could fall between 0.5-2.0, indicating a decline in functional performance spanning areas of memory, orientation, home and hobbies, community affairs, and personal care. For those with a Dementia determination, again a cognitive complaint form self or other was present along with performance across two or more cognitive tests at or below 2 standard deviations from the norm and a CDR SOB greater than or equal to 2.5. All participants for HABS-HD are seen at a single study site at the University of North Texas Health in Fort Worth, Texas. All aspects of the study and study protocol are overseen by the North Texas Regional Institutional Review Board. All HABS-HD participants (and or legal guardians) undergo an informed consent process and provide written authorization for engagement in the study.

We used these racial/ethnic groups as a sensitive attribute. The dataset comprises 4,360 samples and 1,469 features. The missing rate of the original dataset was 31.88%. Initially, we removed each sample and feature that had a *>* 70%missing rate. As the dataset includes multiple visits per individual, we used only the last visit information for each individual in our analysis. However, if any features were missing in the last visit, we imputed those missing values using available data from the same individual’s previous visits. If missing values still remained, we grouped samples based on ethnicity/race and gender to ensure that the nearest neighbors were demographically similar. We then applied K-Nearest Neighbor (KNN) imputation within these groups to fill in the remaining missing values. We used k=3 for KNN imputation in this analysis. However, in rare cases, if all three nearest neighbors of a target individual also had missing values for a specific feature, KNN could not impute those values. For those cases, we subsequently applied an Iterative Imputer. It predicts each feature with missing values using the other available features, processing one feature at a time in a Round Robin manner. Our final dataset had 2,060 individuals and 1,017 features. **Table 1** shows the distribution of HABS-HD samples across the three racial/ethnic groups. In this study, we addressed a multi-class classification problem for AD diagnosis, where the labels comprise three categories: 0 (CN), 1 (Mild Cognitive Impairment), and 2 (AD).

### 4.2 Feature Selection

We implemented three types of feature selection methods: filter-based, wrapper-based, and embedded methods. Below, we explain each type.

#### Filter-Based Methods

Here, we used *Mutual Information (MI)* [21] to measure the association between each feature and the target variable.

#### Wrapper-Based Methods

Wrapper-based methods find the best-performing subset of features using an ML method. For every feature selection method, we used Random Forest (RF) as the base model to predict diagnosis and computed the variable importance score for each feature.

We applied several wrapper-based feature selection methods such as *Recursive Feature Elimination (RFE)* [22], *SelectFromModel* [23], and *SHapley* Additive exPlanations (SHAP) [24]. For example, RFE used the variable importance score to iteratively remove the least important features. SelectBySingleFeaturePerformance, on the other hand, evaluates each feature individually by training a model on it alone and measuring performance.

#### Embedded Methods

We utilized *LASSO* [25] as a built-in feature selection model.

*LASSO* introduces a regularization penalty (controlled by the hyperparameter *λ*) that shrinks coefficients of less important features toward zero, effectively removing those features from the model. We tuned the regularization strength *λ* and found that 0.0003 was the optimal value for *λ* for selecting the feature subset.

### 4.3 Model Training

We trained ML models in four different scenarios to test different hypotheses (**Figure 1**).

**Figure 1:**
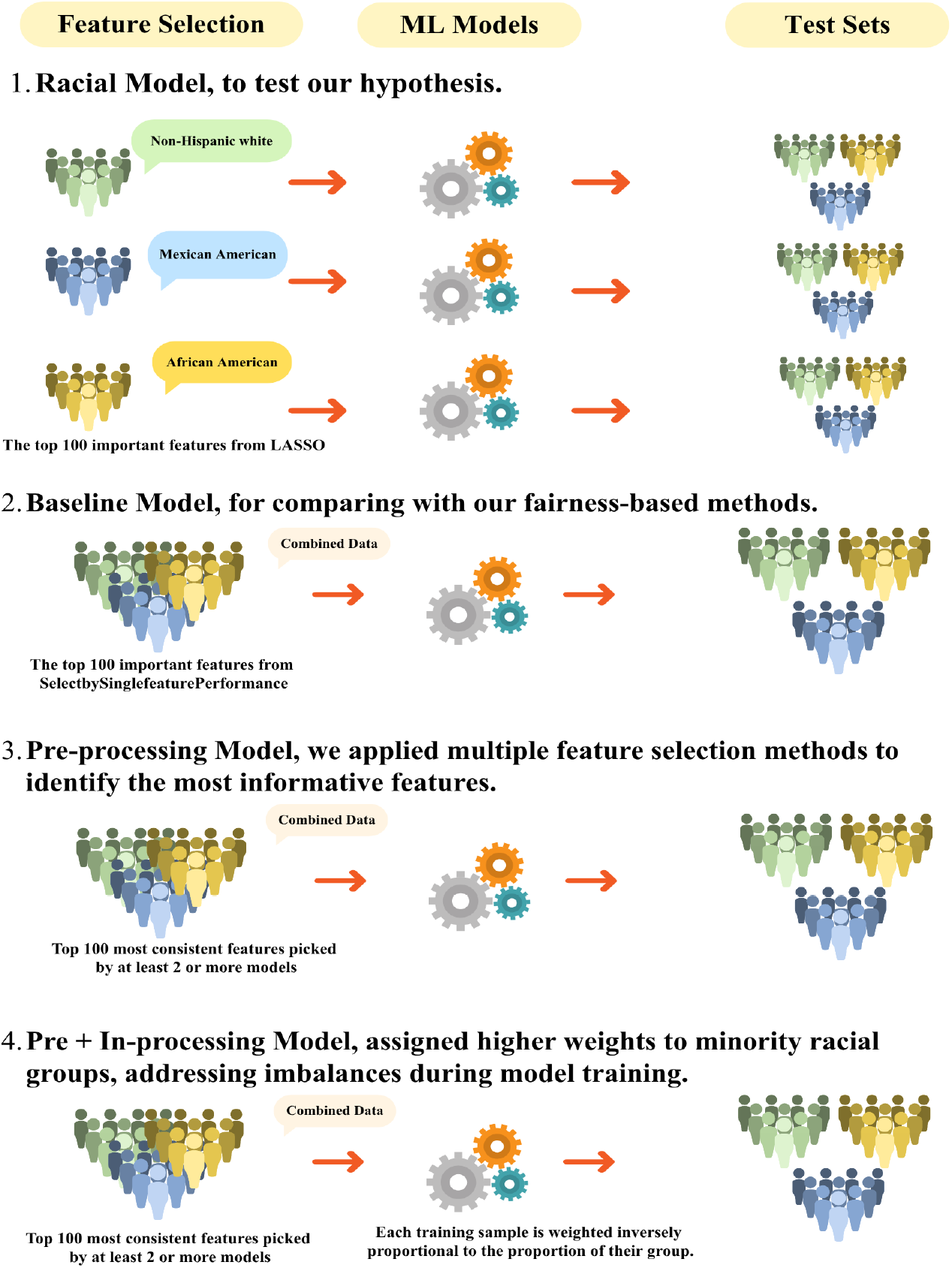
End-to-end pipeline for evaluating and mitigating racial bias in Alzheimer’s prediction. (1) Race-specific model uses top 100 LASSO features, (2) baseline model uses combined data with top 100 SelectbySinglefeaturePerformance features, and (3) Pre-processing model uses 100 most consistent features picked by at least 2 or more models. (4) Pre +In processing model uses 100 most consistent features picked by at least 2 or more models and applies inverse group-proportion reweighting during training, and is evaluated across racial test sets.

First, to test our hypothesis that models trained on one racial/ethnic group can be biased,we performed feature selection using data from a single racial/ethnic group and selected the top 100 important features (giving the best result based on multiple runs), and then evaluated model performance across all racial/ethnic groups (See **Table 2** and **Table 3**). Among the feature selection methods, we used *LASSO* as it demonstrated the best performance in this scenario.

Second, to assess whether training on a racially/ethnically diverse dataset leads to equitable performance across racial/ethnic groups, we trained our models on data consisting of all racial/ethnic groups and tested across different racial/ethnic groups separately. Here, we used the top 100 important features derived from *SelectbySinglefeaturePerformance*, as it showed the best performance compared to the other feature selection methods in this scenario. This model represents the *Baseline* in **Table 4** for comparison to our fairness-based methods.

Third, to mitigate the variability introduced by individual feature selection methods, we employed multiple feature selection techniques to identify the most informative features (See Section 4.2). Within each method, we first performed median-based filtering separately on positive and negative feature importance scores, retaining features with stronger contributions. To harmonize feature importance scores across multiple features, we took the absolute value of each score and applied z-score normalization. Altogether, the methods produced 519 candidate features. To ensure a fair comparison across different settings, we retained the 100 most frequently selected features from this set. This model corresponds to the Pre-processing model in **Table 4**.

Fourth, in addition to the previous scenario, to further increase the fairness of the model, we employed an *in-processing* method where each training sample is weighted inversely proportional to the proportion of its group in the training data. This weighting approach assigns higher weights to minority racial/ethnic groups and underrepresented classes, addressing imbalances during model training. We used the same features used in the third scenario and retrained the model using these weights. This model represents the *Pre-&In-processing* in **Table 4**.

### 4.4 Models and Metrics

Given the class imbalance in our data, we used stratified sampling to divide the data into training and testing sets while maintaining the proportion of class labels and sensitive attributes (i.e., race). Feature normalization was performed after splitting to prevent any data leakage. We used the min-max normalization method, as it maintains the relationships between the original data points, which is important for maintaining the dataset’s integrity. It also helps reduce the impact of outliers by compressing the data range. We utilized several ML models, including Logistic Regression (LR), Gaussian Na ïve Bayes (GNB), Support Vector Machines (SVM), K-Nearest Neighbors (KNN), Multi-layer Perceptron (MLP), and RF. Among these, RF consistently achieved the highest performance across evaluation metrics (See **Supplementary Figure 1**). To ensure a fair comparison, we used the RF models across all the above-mentioned scenarios.

We used macro F1 score and balanced accuracy as evaluation metrics to provide a more reliable assessment of model performance. The macro F1 score is the unweighted average of F1 scores computed for each class individually, ensuring that all classes, regardless of sample size, contribute equally to the final score. Balanced accuracy is calculated as the average of recall (sensitivity) scores across all classes, as these metrics provide a more comprehensive view of how well the model performs across all classes.

### 4.5 Extension of Fairness Metrics to Multiclass Classification

Most of the existing fairness metrics for ML models are primarily designed for binary classification settings [26]. However, real-world datasets often involve multiclass targets, necessitating an extension of these fairness metrics. In our work, we adapted and evaluated three widely recognized group fairness metrics, *Demographic Parity (DP), Equal Opportunity (EOp)*, and *Equalized Odds Difference (EOd)* for a multiclass classification problem with three classes and across three sensitive groups (i.e., AA, MA, and NHW). We define our metrics based on the definition from [27].

#### Demographic Parity (DP)

DP requires that the predicted outcome be statistically independent of the sensitive attribute (e.g., race and gender). In a multiclass setting, this is extended by ensuring that the probability of being assigned to each class is the same across groups. For each racial/ethnic group *r* ∈*R* = {African American, Mexican American, Non-Hispanic White}, we computed the positive prediction rates (PPR) across each class *c* ∈*C* = {0, 1, 2}, *PPR*_*r,c*_ as:

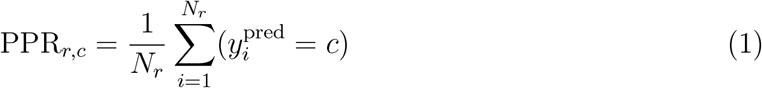

where *N*_*r*_ denotes the number of samples in racial/ethnic group *r*. The DP was calculated by taking the Maximum of the Absolute Difference (MAD) of PPR across the racial/ethnic groups for each class.

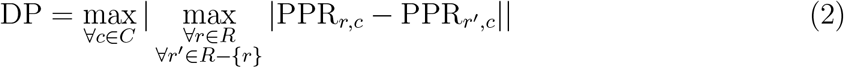

where, *r*′indicates all racial/ethnic groups other than *r*.

#### Equalized Odds Difference (EOd)

EOd requires that both the True Positive Rate (TPR) and the False Positive Rate (FPR) be equal across groups for each actual class label.

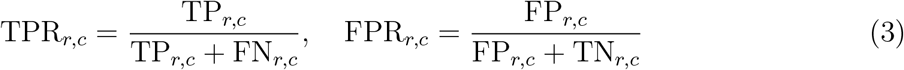

where TP_*r,c*_, FP_*r,c*_, FN_*r,c*_, and TN_*r,c*_ denote the number of true positives, false positives, false negatives, and true negatives, respectively, for group *r* and class *c*.

Similar to Eq. 2, we get the maximum of the Absolute Difference of TPR and FPR rate across the racial/ethnic groups for each class as follows:

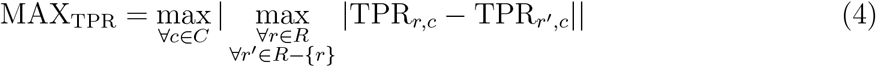

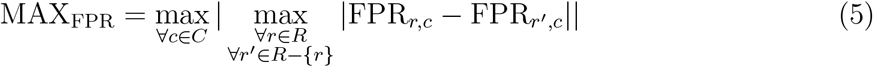

The EOd is calculated as follows:

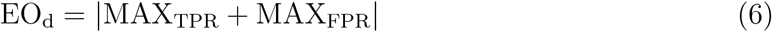

#### Equal Opportunity (EOp)

EOp is a relaxation of EOd that focuses only on the True Positive Rate. It ensures that individuals who belong to class *c* have equal chances of being correctly classified, regardless of their group. This metric is particularly important in high-stakes decision-making contexts (e.g., healthcare and hiring), where true positives represent beneficial outcomes. EOp is calculated using Eq. 4

## DATA AVAILABILITY

The HABS-HD dataset analyzed in this study is available from the HABS-HD study investigators, subject to data access approval and applicable data use agreements.

## CODE AVAILABILITY

The code used for data preprocessing, model training, feature selection, statistical testing, and fairness analysis is available on https://github.com/bozdaglab/AD_ML_Fairness_Evaluation.

## ACKNOWLEDGEMENTS

This work was supported by the National Institute of General Medical Sciences of the National Institutes of Health under Award Number R35GM133657. We thank the HABLE study for the HABS-HD Dataset, Research reported in this publication was supported by the National Institute on Aging of the National Institutes of Health under Award Numbers R01AG054073 and R01AG058533, R01AG070862, P41EB015922, and U19AG078109. The content is solely the responsibility of the authors and does not necessarily represent the official views of the National Institutes of Health.

## AUTHOR CONTRIBUTIONS

Neha Goud Baddam conceived the study, designed the methodology, performed data pre-processing, implemented machine learning models, conducted experiments, analyzed results, and drafted the manuscript. Bizhan Alipour Pijani contributed to the development and implementation of feature selection methods and assisted in result interpretation. Melissa Petersen provided domain expertise on the HABS-HD dataset, contributed to data interpretation, and reviewed the manuscript. Serdar Bozdag supervised the project, contributed to the study design, provided guidance on methodology and analysis, and critically reviewed the manuscript. All authors reviewed and approved the final manuscript.

