## Supplementary for "Evaluating Fairness and Generalizability of Alzheimer’s Disease Diagnosis Models Trained on Racially Imbalanced Datasets"

### Supplementary Materials

#### Supplemental Figure 1

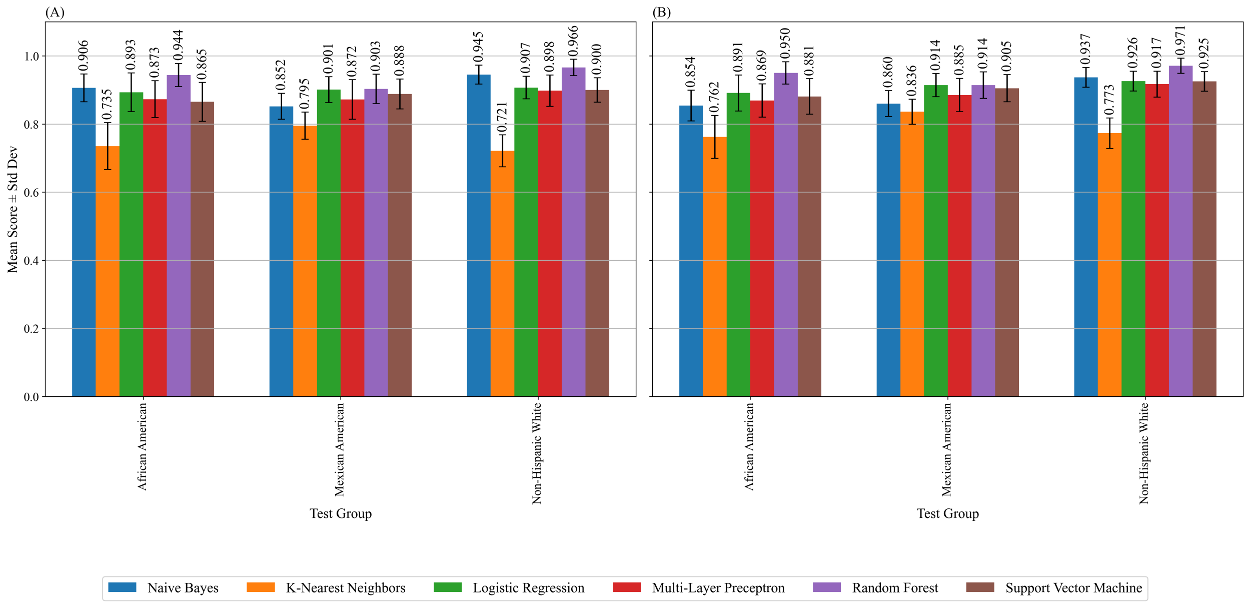


**Supplemental Figure 1.** Performance metrics of different models when trained on samples including all racial groups and separately tested across each racial group. (A) Balanced Accuracy. (B) Macro F1 score.
